# Whole Transcriptome and Functional Analyses Identify Novel Genes Involved in Meiosis and Fertility in *Drosophila melanogaster*

**DOI:** 10.1101/2023.05.12.540472

**Authors:** Siqi Sun, Tyler Defosse, Ayla Boyd, Joel Sop, Faith Verderose, Diya Surray, Mark Aziz, Margaret Howland, Siwen Wu, Neha Changela, Janet Jang, Karen Schindler, Jinchuan Xing, Kim S. McKim

## Abstract

Reproductive success requires the development of viable oocytes and the accurate segregation of chromosomes during meiosis. Failure to segregate chromosomes properly can lead to infertility, miscarriages, or developmental disorders. A variety of factors contribute to accurate chromosome segregation and oocyte development, such as spindle assembly and sister chromatid cohesion. However, many proteins required for meiosis remain unknown. In this study, we aimed to identify and characterize novel meiotic and fertility genes using the genome of *Drosophila melanogaster*. To accomplish this goal, genes upregulated within meiotically active tissues were identified. About 200 genes with no known function were silenced using RNA interference (RNAi), and the effects on meiosis and fertility were assessed. We identified 65 genes that when silenced caused infertility and/or high levels of chromosomal nondisjunction. The vast majority of these genes have human and mouse homologs that are also poorly studied. Through this screening process, we identified novel genes that are crucial for meiosis and oocyte development but have not been extensively studied in human or model organisms. Understanding the function of these genes will be an important step towards the understanding of their biological significance during reproduction.

**Author Summary:** In this study, we aimed to identify and characterize novel meiotic and fertility genes within the genome of *Drosophila melanogaster*. We identified 65 genes that when silenced caused infertility and/or high levels of chromosomal nondisjunction. The vast majority of these genes have human and mouse homologs that are also poorly studied. Through this screening process, we identified novel genes that are crucial for meiosis and oocyte development, making them strong candidates for future studies to characterize their functions.

## Introduction

Fertility requires both a successful meiosis to provide balanced genetic complement to offspring, and several developmental processes to make viable zygotes. During meiosis, germ line cells undergo a single round of genome duplication followed by two consecutive chromosomal divisions prior to fertilization. The meiotic process is highly regulated by multiple cellular structures and protein complexes, but the processes are error-prone, especially in oogenesis. In humans, chromosome segregation errors during oocyte meiosis increase with age. This increase could be related to the unique features of oocytes, such as the extended meiotic arrest, the absence of centrosomes, and possibly other endogenous and exogenous factors [1]. Failure to produce high-quality gametes leads to infertility, spontaneous abortions, and birth defects [2]. Many genes have been identified that are responsible for accurate meiotic divisions and regulate discrete cell-cycle phases, meiotic events, and gamete viability [3–6]. Nevertheless, we still lack a complete picture of the proteins that control key processes in meiosis, such as chromosome cohesion, chromosome biorientation, and spindle assembly.

In addition to meiosis, defects in numerous developmental processes can also lead to infertility, such as genes required for germline development [7]. Furthermore, in most animals, maternally derived gene products regulate the early events of embryogenesis [8]. These include genes required for egg activation, early mitotic divisions, and the maternal-to-zygotic transition [9–11]. Elimination of these genes in the mother can lead to loss of fertility. It is likely that many additional factors are responsible for the loss of female fertility in humans, although few have been identified [5, 12].

*Drosophila* is a simple but vital genetic model for identifying and understanding the function of genes required for germline and embryonic development and meiosis [13, 14]. Despite the evident differences between *Drosophila* and human physiology, the homologs of many human genetic disease loci show selective expression in *Drosophila* tissues analogous to the affected human tissues [15]. Additionally, the ability to produce a large number of offspring in a short period of time makes *Drosophila* a powerful system for gene discovery and studying potential disease-causing genes [16, 17].

In *Drosophila*, many well-studied genes required for meiosis are upregulated within the ovary, such as components of the synaptonemal complex *c(2)M* and *c(3)G* [18, 19] and members of the Chromosome Passenger Complex such as INCENP, and Aurora B kinase [20]. Genes that are required for oocyte and early embryonic development are also upregulated in the ovary, such as *nanos*, *vasa*, and *bicoid* [21–24]. By considering a gene’s expression pattern, sequence homology, and protein folding patterns, it is possible to predict the function and subcellular localization of novel proteins. Such information would allow prioritization of screens based on the likelihood of a gene being involved in a biological process such as meiosis. Several online databases provide tissue-specific expression profiles and functional annotations of *Drosophila* genome, and they could be used to identify potential genes required for meiosis and fertility [15, 25–27]. In this study, we identified and validated 65 novel genes involved in meiosis and fertility by combining tissue-expression profile, gene annotation, functional analysis with RNAi knockdown, and cytological analysis. When these genes were knocked down in the germline, the flies displayed meiosis or germline development related phenotypes, including sterility, reduced fertility, and elevated levels of nondisjunction.

## Methods

### Candidate gene selection from FlyAtlas 1

Gene expression profiles in different fly tissues were downloaded from FlyAtlas 1 (http://flyatlas.org/data.html) [15]. A total of 18,770 probe sets from 13,500 genes were included on the Affymetrix *Drosophila* Genome 2 expression array. The gene names and Flybase IDs were validated and corrected by FlyBase (https://flybase.org/). Genes with the following symbols in their names were removed, including unknown genes (“---”), non-protein-coding genes (“CR”), RNA genes (“rna”), and ribosomal proteins (“rps”/“rpl”). Expressions in tissue “*Drosophila* S2 cells” were excluded from analysis.

For each gene, the enrichment value for each tissue was calculated as the tissue-specific mean expression divided by the mean of the fly whole body; the p value was calculated using two-tailed Student’s t test. The expression direction is categorized as “up” when the enrichment value is larger than 1 and the p value is less than or equal to 0.05 (enrichment > 1 and p-value <= 0.05), “down” as enrichment < 1 and p-value <= 0.05, and “others” for anything else. To select genes with high confidence, microarray probe sets with no signals detected among four biological replicates (i.e., no “present” calls) in ovary, testis, and larvae central nervous system (CNS) were removed. One representative probe set was selected for each gene by selecting the one with the most biological replicate results and direction being “up” in ovary, testis, and larvae CNS. The selected genes were divided in four groups: ovary_only (“up” in ovary expression and not “up” in other tissues), ovary+cns (“up” in ovary and larvae CNS expression, not “up” in other tissues), ovary+testis (“up” in ovary and testis expression, not “up” in other tissues), and ovary+CNS+testis (“up” in ovary, larvae CNS, and testis expression, not “up” in other tissues).

### Candidate gene annotation and prioritization for experimental validation

To identify novel meiosis gene for functional validation, gene symbols with the format “CG + numbers” were considered to be not-well studied and selected. Next, the transgeneic RNA interference line for the candidate genes were searched in the Transgenic RNAi Project (TRiP) [28, 29] at Bloomington Drosophila Stock Center (https://bdsc.indiana.edu/index.html). Candidate genes without TRiP stocks were removed.

Experimentally-validated and predicted meiosis genes in *Drosophila* were extracted from MeiosisOnline (https://mcg.ustc.edu.cn/bsc/meiosis/index.html) [30]. Gene expression value of three ovary cell clusters, germ cells (GC), germarium soma, and follicle cells (FC), were calculated using single-cell transcriptome data [31]. For each cluster, the expression level of each gene was calculated as the median value of expressions from individual stages as defined in the Supplemental Table S2 of [31].

### Candidate gene annotation, enrichment and protein-protein interaction network analyses

Orthologs and known alleles of candidate genes were annotated using FlyBase [27]. Enrichment analyses were performed for positive genes with human and mouse orthologs using the overrepresentation analysis in ConsensusPathDB (CPDB, http://cpdb.molgen.mpg.de/) [32]. Enriched terms (e.g., Gene Ontology, pathway, or protein complex) containing at least two input genes were selected for further analysis. Enrichment p values were determined by CPDB using a hypergeometric test and q values represent the adjusted p values using the false discovery rate method.

Protein-protein interaction (PPI) network of positive genes was constructed using interactions information provided by STRING *Drosophila melanogaster* database [33]. Interactions with at least medium confidence (i.e., combined scores >= 0.4) were selected.

### RNAi knockdown and sterility and nondisjunction assays

Potential meiotic genes were screened using RNA interference (RNAi). Stocks for RNAi were obtained from the Bloomington Drosophila Stock Center at the Indiana University Bloomington (https://bdsc.indiana.edu/index.html). In these transgenes, the shRNA sequence is placed downstream of the UAS enhancer, thus requiring the presence of the GAL4 activator to begin gene knockdown via RNAi.

Crosses were set up using 10-20 *UAS:shRNA* males and 15-25 females with a tissue-specific *GAL4*. Each shRNA was crossed to two *GAL4* stocks to induce expression. *P{GAL4: :VP16-nos.UTR}CG6325^MVD1^* (referred to as *MVD1*) initiates expression of transgenes in the pre-meiotic mitotic cells of the germline and continues throughout oocyte development [34]. *P{w[+mC]=matalpha4-GAL-VP16}V37* (referred to as *matα*) expression begins late in prophase (region 2b/3), after early pachytene and the formation of crossovers and the synaptonemal complex, and continues through late prophase until stage 14 oocyte [20, 35].

All crosses were kept at 25°C and were allowed to incubate for 10 days until progeny began to emerge. Virgin female progeny from this cross that carried both the UAS RNAi and the *GAL4*, as indicated by the y+ and w+ phenotypes, were collected. Five *UAS RNAi/GAL4* females were then crossed to five *y w /Y, B^s^* males in five sets of vials (replicates). On days 14 and 18, the crosses were scored by recording the number of wildtype females (X/X), Bar males (X/B^S^ Y), aneuploid males (X/O), and aneuploid females (X/X/B^S^ Y). The frequency of NDJ was calculated as 2(XO + XXY) / [2(XO + XXY) + X/X + X/Y]. An RNAi line is considered positive if the cross has at least one of the following phenotypes: 1) sterile; 2) NDJ frequency >= 1.7% in either *MVD1* or *matα*, or 3) <=100 progeny in either *MVD1* or *matα*. A candidate gene is considered a positive gene if at least one of its shRNA lines is positive. Some positives from these two tests were crossed to *P{w[+mC]=tubP-GAL4}LL7* (referred to as *Tub:GAL4*), which expresses throughout the whole body of the fly and is a test for a somatic function such as mitosis.

### Cytology

To study the effects of target gene knockdowns on germline development, meiosis, and embryonic development, the ovaries were examined. Small ovaries are indicative of a failure in germline development, such as loss of germline stem cells or development of the ovarian cysts.

For cytological examination of early (germarium/ pachytene) or late (stage 14/ Metaphase I) meiosis, oocytes were collected and examined using immunofluorescence. Crosses were set up using 10-20 *UAS:shRNA* males and 15-25 tissue-specific *GAL4* females. Twelve days following the cross, approximately 20 (for germarium) or ~300 (for stage 14) progeny that carried both the UAS regulated RNAi (y+) and the tissue-specific *GAL4* (w+) were collected and fed yeast for 2 days at 25°C to promote egg laying. After 2 days, oocytes were collected and fixed using whole mounts to preserve the three-dimensional structure. Details of the two fixation protocols are described elsewhere [36, 37].

The primary antibodies used were: mouse anti-C(3)G (1:500) [38], rabbit anti-C(2)M (1:400) [18], a combination of two mouse anti-ORB antibodies (4H8 and 6H4, 1:100) [39], rabbit anti ᵧH2AV (1:500) [40], mouse anti-α-tubulin conjugated to FITC (1:50) to stain microtubules, rabbit anti-CID (1:1000), and rat anti-INCENP (1:600). Additionally, Hoechst 33342 was used to stain for DNA. Following overnight incubation, oocytes were washed and stained with secondary antibodies for 4 hours at room temperature. The secondary antibodies used were: goat anti-rat Cy3 (1:100), goat anti-rabbit 647 (1:100), and Alexa 488 (1:200) (Jackson Labs and Invitrogen). Oocytes were then mounted in SlowFade Gold (Invitrogen) and imaged using a Leica TCS SP8 confocal microscope with a 63x, NA 1.4 lens. All images shown are maximum projections of complete image stacks. Statistical analysis of sister kinetochore separation and foci quantification were performed using the microscopy image analysis software Imaris (Oxford Instruments).

## Results

### Selection of genes up-regulated in the ovary

FlyAtlas 1 contains expression levels of 13,500 genes in *Drosophila* adult and larval tissues. We selected 10,948 genes that are expressed in ovary, testis, and larval CNS. Genes that were up-regulated in adult ovaries while not up-regulated in other tissues except testis and larval CNS were selected as initial genes of interest (see Methods for detail). This selection resulted in 1,119 genes, 611 of which were up-regulated only in ovary (referred to as “ovary-specific” in the following text). The ovary-specific genes included known meiotic genes, such as *c(2)M* and *c(3)G.* Among the 1,119 genes, 141 were up-regulated in both ovary and testis. These genes were included because testis is a meiotic tissue. Genes in this selection included *ord,* a gene required for sister chromatid cohesion. Finally, because of mechanistic conservation in segregating chromosomes, we included genes required for meiosis that are also required for mitosis. Therefore, we included genes up-regulated in ovary and larval CNS because the larval CNS is a mitotically active tissue. Among genes upregulated in ovary, 306 were also up-regulated in CNS and 61 were up-regulated in both CNS and testis (Figure 1). Examples of these genes included spindle-associated proteins such as the CPC components *Incenp* and aurora kinase B (*aurB*), and kinetochore proteins such as *Ndc80* and *Spc105R.* Among the 1,119 genes, 975 have human orthologs and 965 have mouse orthologs. Importantly, 874 of these genes have a shRNA TRiP stock which makes phenotype screening feasible [28] (Figure 1, Table S1). To identify potential novel meiosis genes, we selected a subset of uncharacterized genes, many of which lack a gene name, and for whom a shRNA TRiP stock was available. This group was enriched for genes that have not been studied before and have no known function. After further manual review, we selected 205 genes for functional test by RNAi knockdown (Figure 1).

**Figure 1:**
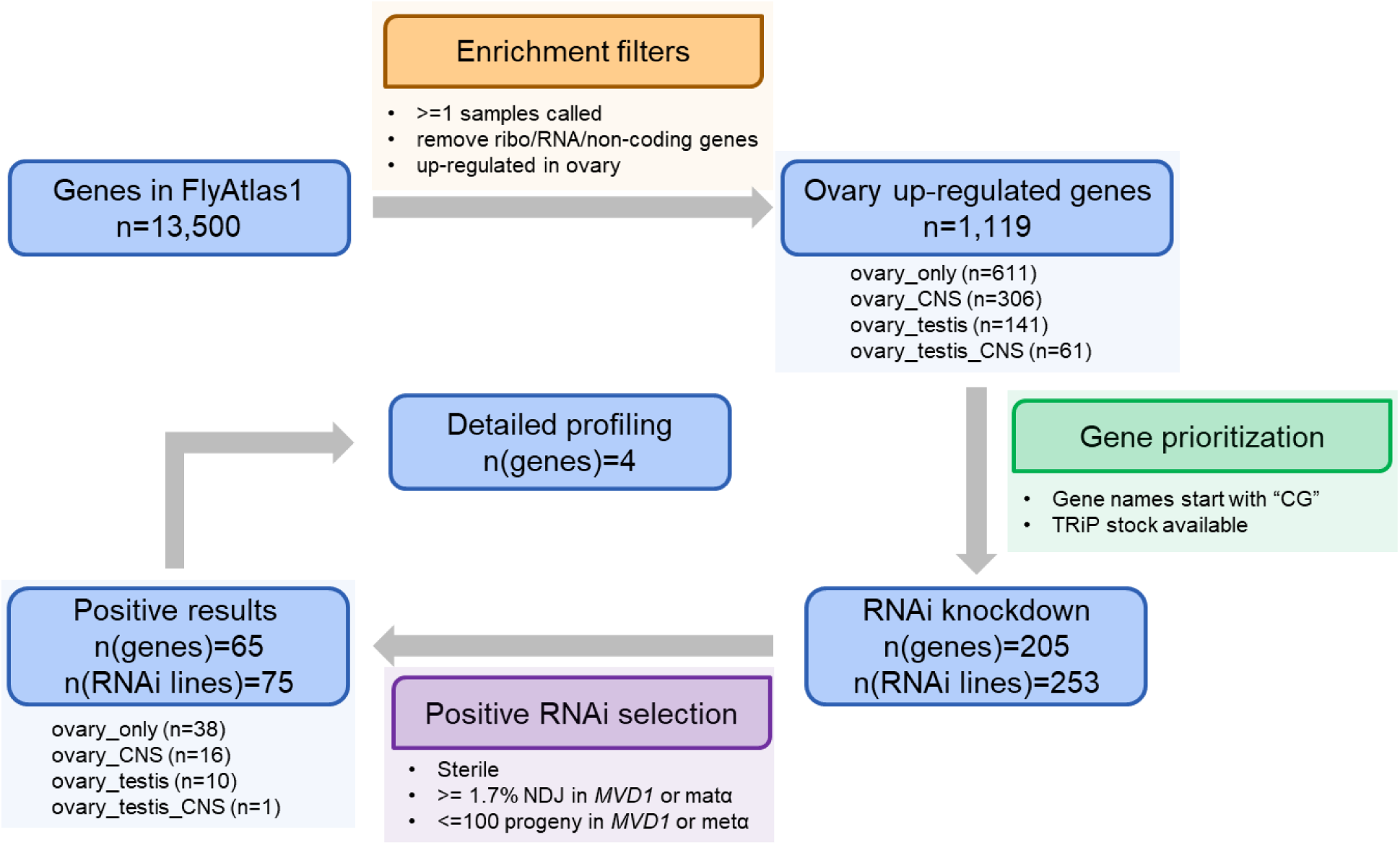
Analysis workflow. The original 13,500 genes in Flyatlas1 were first selected by different enrichment and prioritization filters. The passed genes were validated by RNAi knockdown and sterility and nondisjunction assays. A total of 65 genes and the associated 75 RNAi lines were identified as positive novel meiosis genes with 4 genes further profiled in detail. The numbers of genes in different sub-categories (e.g., ovary_only) were counted. The numbers of genes (n(genes)) and RNAi lines (n(RNAi lines)) after each filter were denoted.

### Ovary-specific RNAi knockdown of candidate genes

Each *shRNA* strain was crossed to one of two GAL4 expressing females to induce shRNA expression. *P{GAL4: :VP16-nos.UTR}CG6325^MVD1^* (referred to as *MVD1*) initiates expression of transgenes specifically in the pre-meiotic cyst cells of the germline and continues throughout oocyte maturation [34]. *P{w[+mC]=matalpha4-GAL-VP16}V37* (referred to as *matα*) expression begins in prophase (region2b/3) and continues through until Metaphase I in stage 14 oocytes [20, 35]. A significant phenotype (a “positive”) was scored if the RNAi knockdown produced sterile females, or an increased nondisjunction rate (>= 1.7%) that was higher than most of RNAi lines (**Figure S 1**), or a small brood size (< 100). Because each gene was tested with *MVD1* and/or *matα* promoters, and some genes were tested using multiple shRNA lines, we defined a positive gene as one with as at least one RNAi experiment yielding a significant result with either the *MVD1* or the *matα* promoter.

Following completion of nondisjunction and sterility assays on 205 novel gene candidates in 253 RNAi lines (Table S2), we identified 65 genes that showed evidence of a function in germline development, meiosis, or embryonic development in experiments with 75 RNAi lines (Table 1, Figure 2). Some positive RNAi lines (e.g., GL00570) are present in multiple phenotype categories (NDJ, Sterile, and small brood size) because: 1) different phenotypes were observed between the MVD1 and the *matα* experiments; 2) phenotypes of some shRNAs met the criteria of both NDJ and small brood size (Figure 2A). Among the 65 genes, 38 are ovary-specific genes, 16 are expressed in both ovary and larval CNS, 10 are expressed in both ovary and testis, and 1 is expressed in all three tissues. Knockdown of 21 genes produced sterile females, 22 genes had an increased nondisjunction rate (>= 1.7%), and 7 genes produced small brood sizes (< 100) (Figure 2B). The remaining 15 genes showed different phenotypes in *MVD1* and *matα* crosses and/or with different shRNAs (overlapping sections of the Venn diagram, Figure 2B). For example, CG10336 *MVD1/* shRNA knockdown females were sterile, but had a small brood size in *matα/* shRNA females. The difference between the two *GAL4* lines provides temporal information on when these genes function during oocyte development.

**Figure 2:**
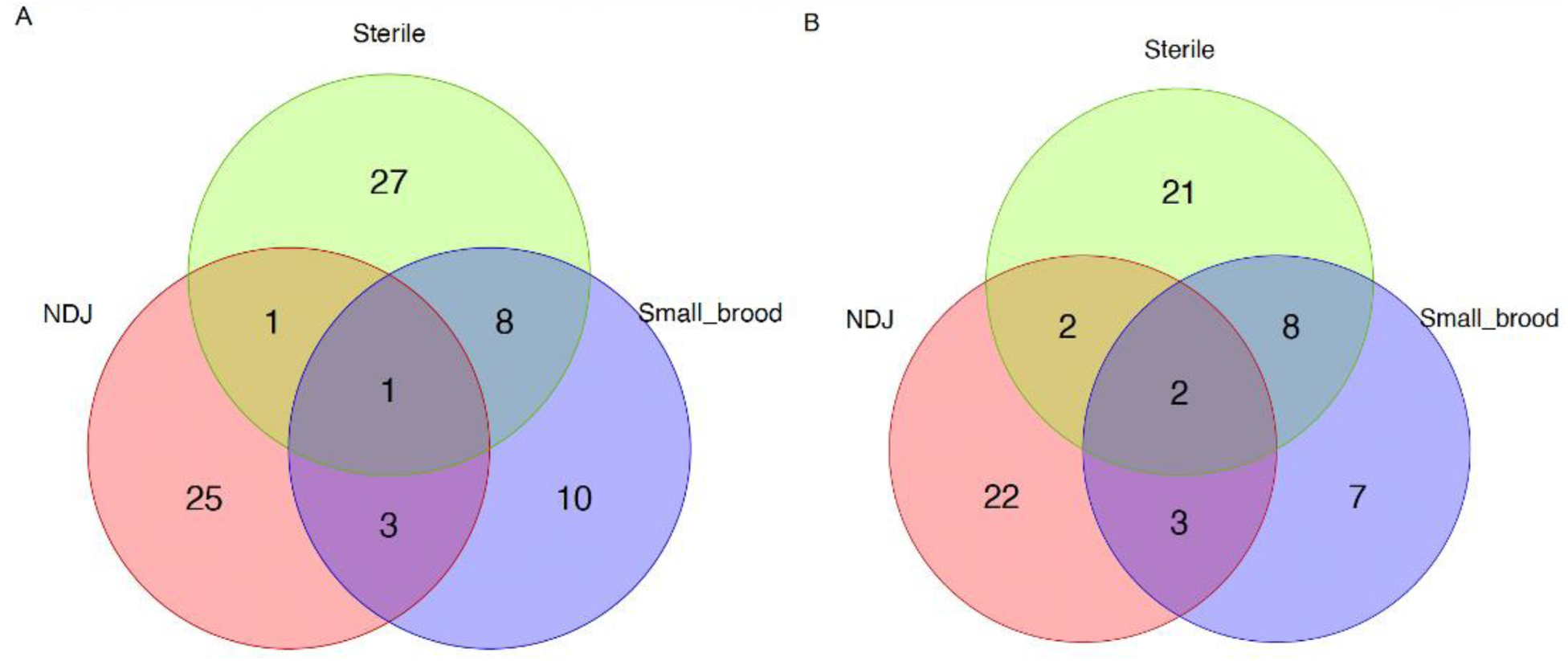
Venn diagrams for (A) 75 positive RNAi lines and (B) 65 positive genes. Some RNAi lines and associated positive genes are present in multiple phenotype categories (NDJ, Sterile, and small brood size (Small_brood)) because: 1) different phenotypes were observed in the *MVD1* and *matα* experiments; 2) phenotypes of some lines met the criteria of both NDJ and small brood size. See details in Table 1.

**Table 1:**
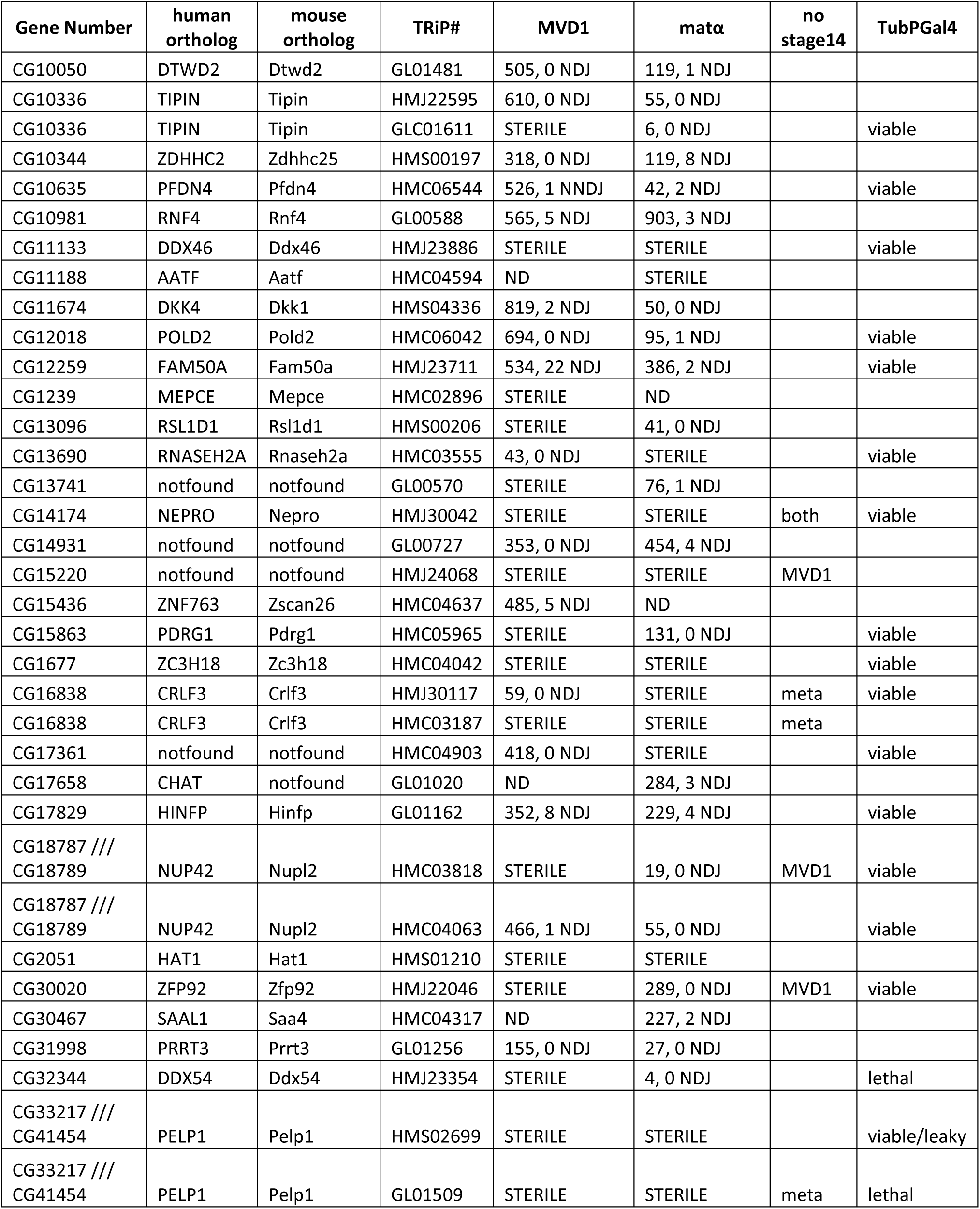
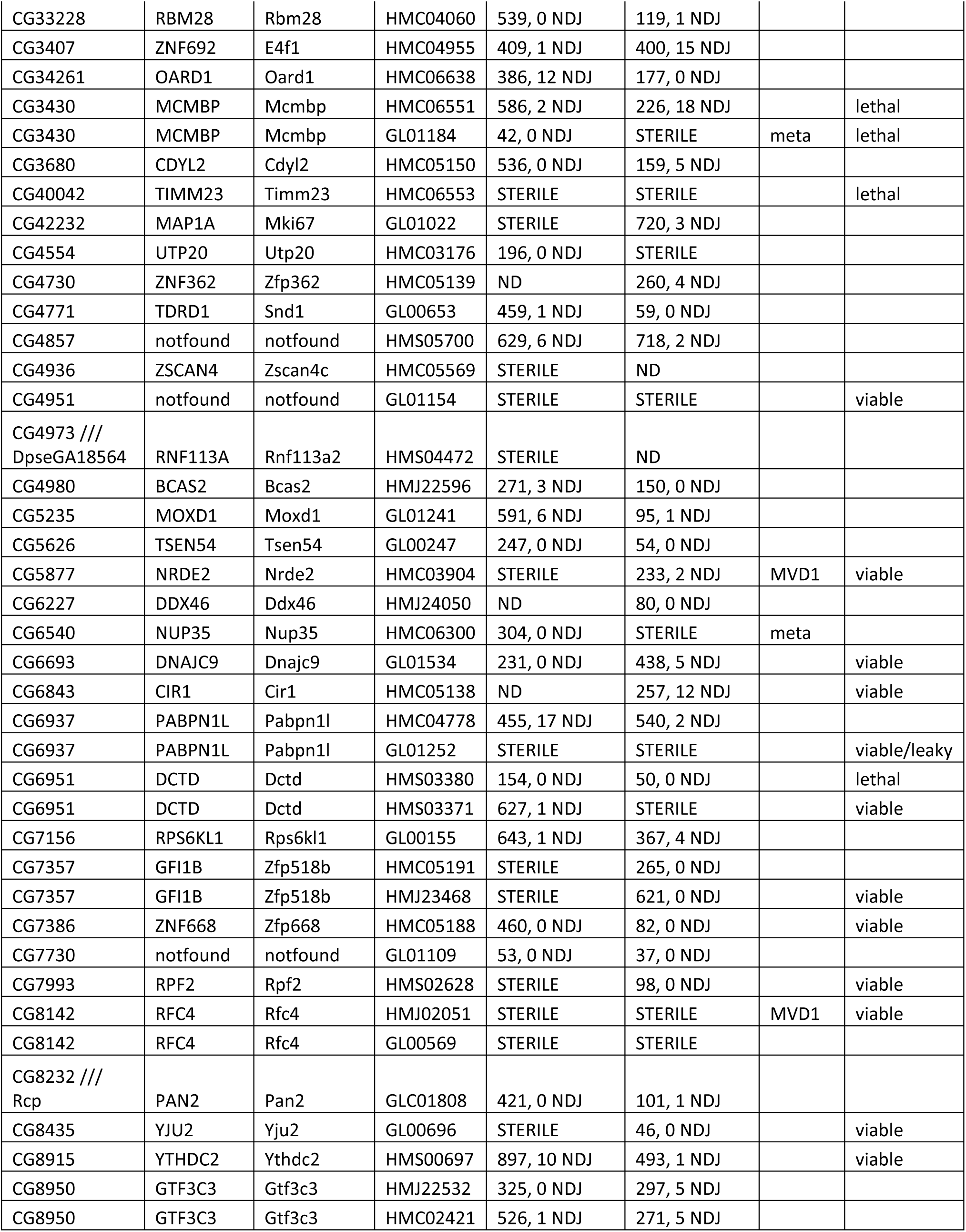
65 meiosis-related genes RNAi knockdown phenotypes. See Table S2 for a full list of tested RNAi lines and genes.

### Annotation and enrichment analyses of positive genes

To identify genes with a role in meiosis, we examined proposed functions of each identified gene in other species (Table 1). Among the 65 genes, 58 and 57 genes have human or mouse orthologs, respectively. Using human or mouse orthologs of the *Drosophila* genes, we performed gene ontology (GO) and pathway enrichment analyses (Table S3, Table S4). Several GO terms showed significant enrichment in human and mouse orthologs, such as replication fork (q<2.4e^-4^ in human and mouse) and DNA polymerase complex (q<1.9e^-3^ in mouse, q<5.3e^-3^ in human, Table S3). Similarly, several pathways showed significant enrichment, such as DNA replication (q<3.1e^-5^ in mouse, q=0.011 in human), DNA repair (q=0.018 in human), and mitotic cell cycle (q=0.013 in mouse, Table S4). These GO terms and pathways might be expected as important for various steps of meiosis or the mitotic divisions of the early embryo. In addition, genes associated with RNA biology, such as spliceosome complex, RNA helicase activity, and ribosome biogenesis, also appeared frequently in the analysis. These findings are consistent with the known importance of regulating RNAs in oocyte and early embryonic development [41–43].

Furthermore, we examined the interaction among the candidate genes by constructing a protein-protein interaction (PPI) network. We identified a network of 37 positive genes in 3 main clusters (Table S5, Figure 3). The clusters contain genes in two significantly enriched GO terms related to cellular component biogenesis and mRNA splicing: ribonucleoprotein complex biogenesis (q=0.0051, CG32344, CG4554, CG13096, CG11188 and CG7993) and spliceosome complex (q=0.0096, CG8435, CG4980, CG4973). These genes could be required for generating oocytes with enough maternal products to support embryonic development. We also found three genes in the GO term HDR through Homologous Recombination (HRR) or Single Strand Annealing (SSA) genes (CG8142, CG10336, CG12018). These genes could be important for meiosis as homology-directed repair is crucial during meiosis to ensure proper chromosome segregation [6].

**Figure 3:**
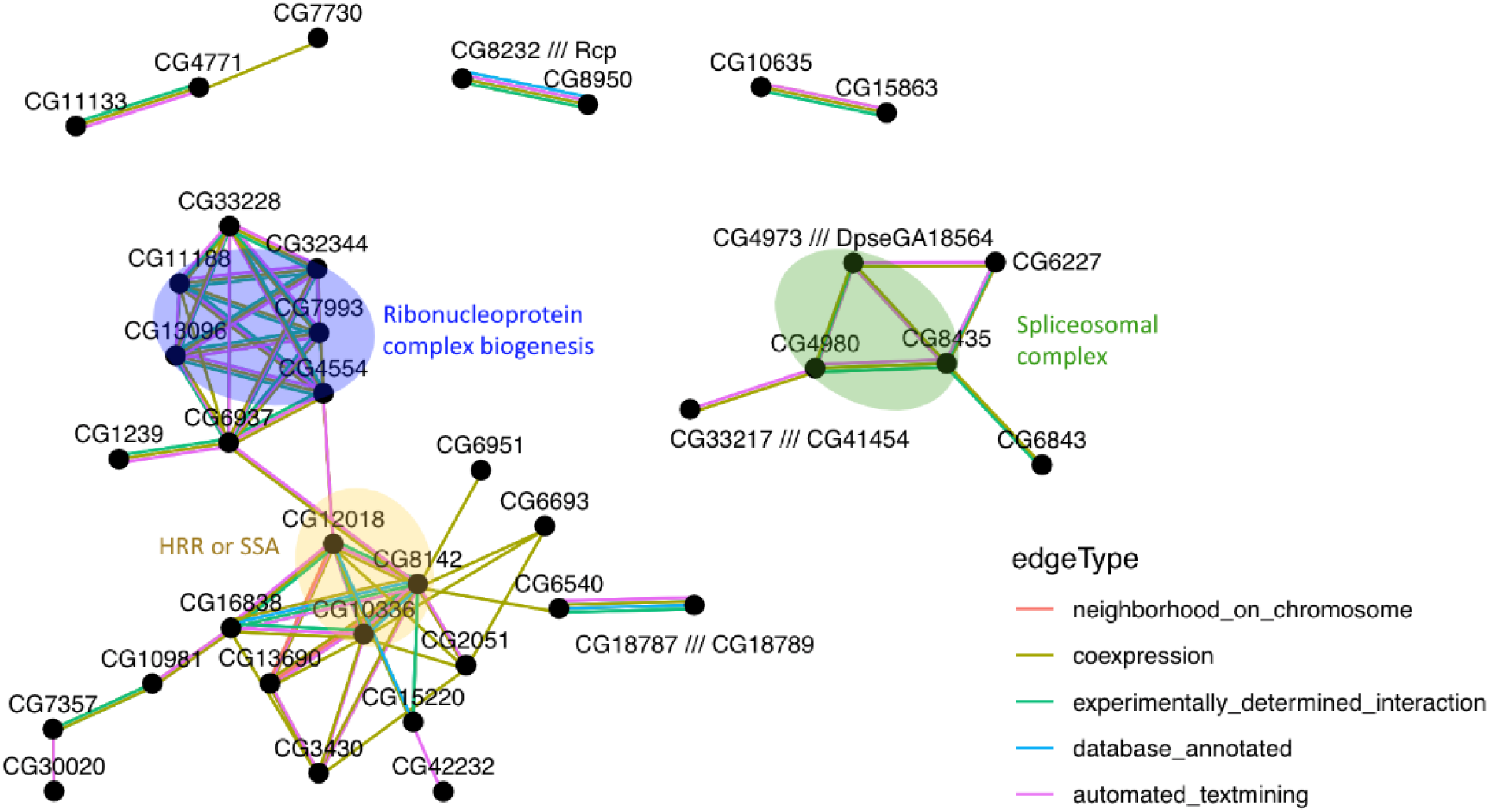
Protein–protein interaction network of 37 candidate genes. Only connected genes are shown. Genes present in three representative enriched GO terms: ribonucleoprotein complex biogenesis, spliceosomal complex, and HDR through homologous recombination (HRR) or single-strand annealing (SSA) are highlighted with the enriched terms labeled. See Supplemental Table S5 for a full list of interactions.

While we selected genes expressed in the ovary, ovaries are a complex tissue consisting of somatic and germline cell types. Insights into the function of these genes could come from identifying in which ovarian cell type they are expressed. A single cell ovary transcriptome dataset has identified several somatic (germarium soma, follicle cell (FC)) and germline cell (GC) cell types in ovaries [31]. Based on this dataset, 908 of the 1,119 ovary up-regulated genes in our initial dataset (85%) had higher expression in the GC cluster compared to the germarium soma and FC clusters, with 84% of positive genes (55 genes) showing the same trend. These results suggest that our candidate genes are highly enriched in germline-specific genes and are likely to function in oocyte development.

### Analysis of ovaries from shRNA lines with fertility or meiotic phenotypes

We selected four genes with a strong genetic phenotype, either sterility or high nondisjunction, for further analysis by cytological screening. Oocytes from shRNAs targeting CG3430, CG4951, CG12259, and CG18787 were examined in early prophase (the germarium) and in mature Metaphase I-arrested oocytes.

### CG3430 knockdown has biorientation defects and precocious anaphase I onset

Two shRNAs were associated with CG3430 (Table 1). The first shRNA, *GL01184*, caused reduced fertility when expressed with *MVD1* and was sterile when expressed with *matα.* The second shRNA, *HMC06551*, had little effect when expressed with *MVD1* and elevated levels of nondisjunction (15.9%) when expressed with *matα*. The differences in these phenotypes indicate that *GL01184* resulted in a stronger knockdown of mRNA than *HMC06551.* Both shRNAs caused lethality when expressed by *Tub:GAL4*, indicating this gene could be required for mitosis as well.

The ovaries of *GL01184/ matα* females lacked stage 14 oocytes, suggesting that *CG3430* is required for oocyte growth or development. To determine if *CG3430* has a role in Metaphase I, stage 14 oocytes from *HMC06551/ matα* flies were examined for cytological defects. In wild-type oocytes, Metaphase I chromosomes are together in a single karyosome at the center of a bipolar spindle (Figure 4A). In addition, biorientation of each kinetochores pair can be observed, such that 4 centromeres, labeled with anti-CID, are directed towards each pole of the Metaphase I oocyte (Figure 4A). In *HMC06551/ matα* females, karyosome separation typical of anaphase was found in approximately 33% (n=15) of Metaphase I oocytes (Figure 4B, 4C). In some of these cases, the karyosome was separated in two distinct masses with a bridge of chromatin between them (Figure 4C). Additionally, we observed evidence of biorientation defects, where kinetochores were asymmetrically distributed in 2/15 oocytes (13%), a phenotype consistent with elevated nondisjunction. To identify defects early in meiotic prophase, ovaries from *GL01184/ MVD1* females were dissected and stained with antibodies that mark Synaptonemal complex (SC) components C(2)M and C(3)G. No cytological defects were observed (compare wild-type in Figure 5A and CG3430 knockdown in 5B), suggesting CG3430 is only required in late meiotic prophase or Metaphase.

**Figure 4:**
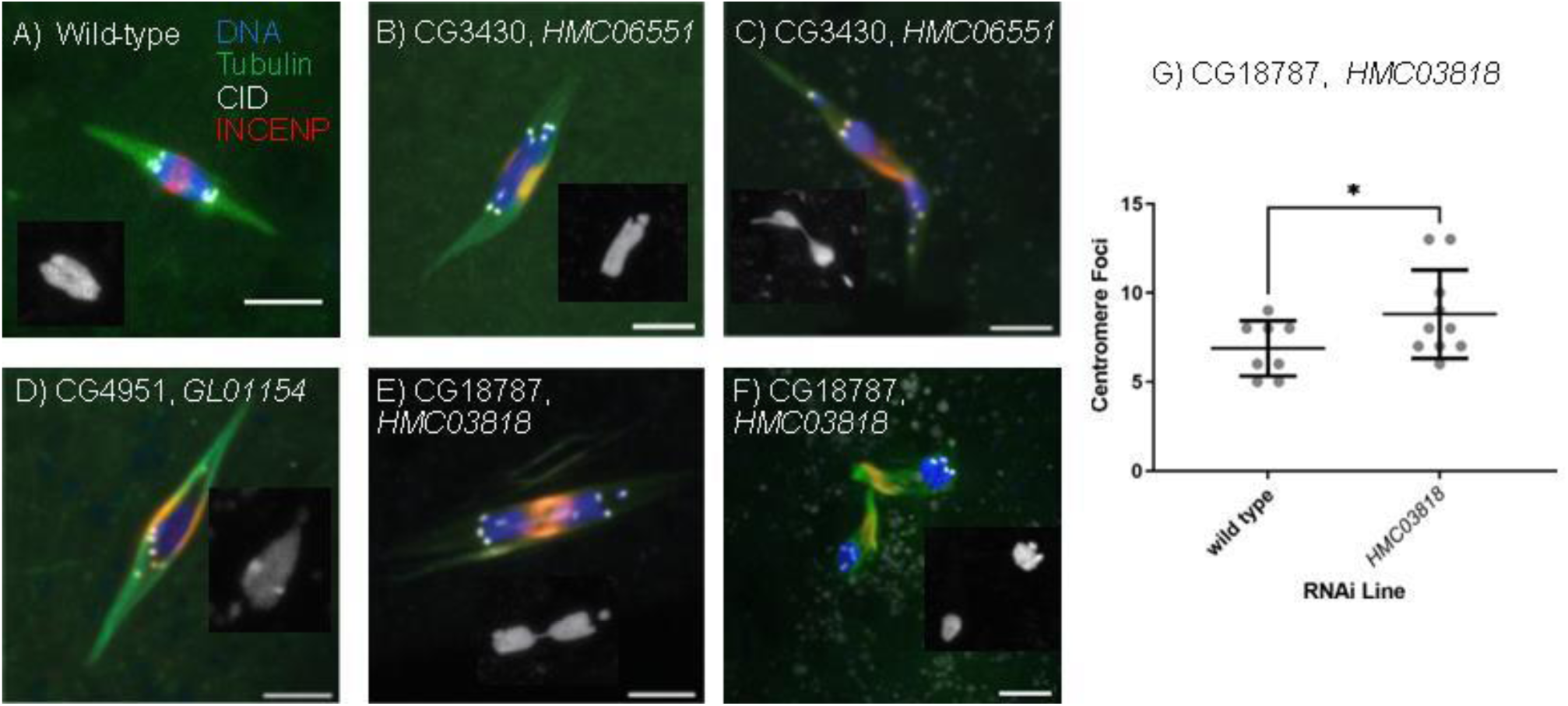
Cytological analysis of *Drosophila* Oocytes. Drosophila oocytes were extracted and stained for Tubulin, the centromere (CID/CENP-A), central spindle (INCENP), and DNA (Hoechst). Scale bars are 5 μm. (A) An example of a wild-type Metaphase I arrested spindle with the formation of a singular karyosome and symmetric division of the centromeres. Knockdown using *matα* of (B,C) CG3430, (D) CG4951 and (E,F) CG18787. Inset in each panel shows the karyosome (DNA). (G) Quantification of sister kinetochore foci for both wildtype and CG18787 RNAi oocytes.

**Figure 5:**
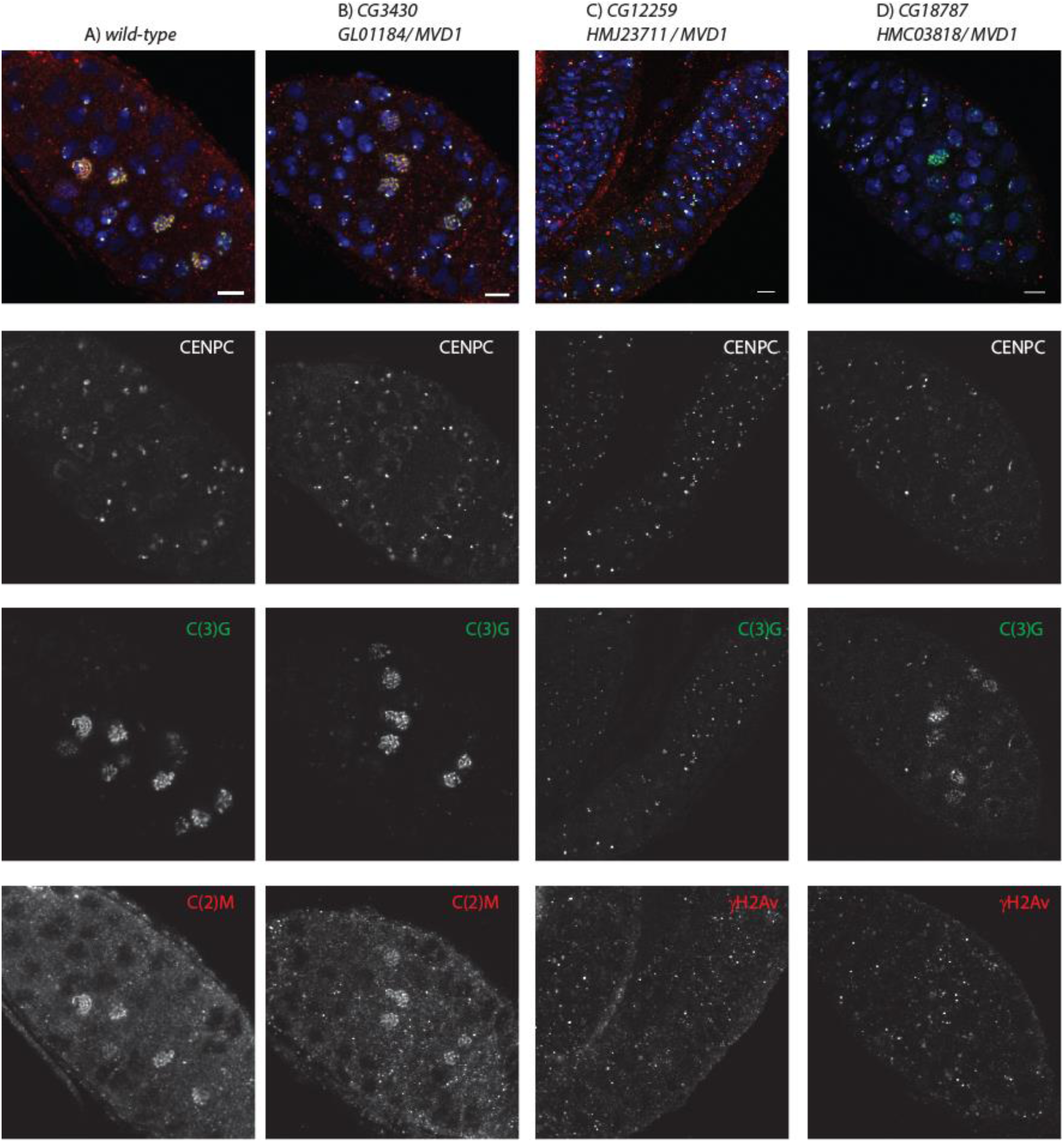
Early prophase (germarium) images. Ovaries were dissected from shRNA / *MVD1* females and stained for C(3)G (green), ᵧH2AV or C(2)M (red), CENP-C (white), and DNA (blue). RNAi against (B) CG3430 and (D) *CG18787* had relatively normal cytology, including development of 16-cell cysts and threads of C(3)G. RNAi against CG12259 (C) had small ovaries characterized by a lack of 16-cell cyst formation and C(3)G limited to the centromeres. This is characteristic of the mitotic (region 1) cells in the germline.

By sequence comparison, we found that CG3430 is homologous to the Mini-Chromosome Maintenance Complex Binding Protein (MCMBP) superfamily. For example, CG3430 has 37% identity with human MCMBP, which was identified as a protein that associates with and promotes assembly of the MCM2-7 complex [44, 45]. Other functions have also been suggested. For example, in *Xenopus*, MCMBP promotes disassembly of MCM complexes from chromatin [46]. *Arabidopsis thaliana* MCMBP/ETG1 appears to be needed for sister chromatid cohesion [47]. This function is consistent with both the nondisjunction and cytological phenotypes observed in this study. Our results with *matα* demonstrate that *Drosophila* MCMBP has a function after premeiotic S-phase in oocytes.

### CG4951 knockdown has biorientation defects

Knockdown of *CG4951* using shRNA *GL01154* caused sterility (Table 1). Sequence analysis of this gene does not provide any significant insight into the function as the predicted protein sequence does not contain identifiable protein domains or regions of high conservation. *CG4951* is an example of a poorly conserved gene. It encodes a 320 amino-acid (aa) protein that is conserved within the *Drosophila* genus but not found in other Diptera such as the mosquito. Cytological analysis of *CG4951* RNAi oocytes revealed evidence of biorientation defects, as shown by asymmetric distribution of kinetochores, which would result in the division of an abnormal number of chromosomes to each pole (Figure 4D, 2/10 oocytes). These results suggest that *CG4951* is involved in the regulation of kinetochore biorientation. *GL01154* did not cause lethality when crossed to *Tub:GAL4*, suggesting the function of *CG4951* is germline specific and not required for mitosis.

### CG12259 knockdown has early defects in germline development

*CG12259* has the highest homology with *FAM50A* in mammals with possible RNA/nucleic acid binding activity and functions in chromatin organization [48]. Two lines of evidence suggest this is a gene required early in germ line development. First, *HMJ23711/ MVD1* females were sterile but *HMJ23711/ matα* females were fertile. Second, cytological analysis of *HMJ23711/ MVD1* females revealed a severe phenotype, with an early defect in germline development and failing to make 16 cell cysts and an oocyte with full SC formation (Figure 5C). Thus, this gene is probably required prior to meiosis, during the mitotic divisions that generate the 16 cell cysts. *HMJ23711* did not cause lethality when crossed to *Tub:GAL4*, indicating *CG12259* may not be required for mitosis.

### CG18787 knockdown has premature karyosome and sister centromere separation

One shRNA targeting *CG18787* could also target a second gene, *CG18789*. These two genes are located in close proximity to one another, separated by approximately 2.5kb and two other genes. *CG18787* encodes a 398 aa protein that has 99% amino acid identity with *CG18789*, with only 3 amino acid differences. Given both the high similarity scores and the relative location of these two genes within the genome, it is likely that *CG18787* and *CG18789* arose from a recent gene duplication event. ShRNA *HMC03818* produced a more severe fertility defect when crossed with both the *MVD1* and *matα* compared to shRNA *HMC04063* (Table 1). Analysis of the sequences of each shRNA demonstrated that *HMC03818* could target both *CG18787* and *CG18789*, while *HMC04063* probably targets only *CG18787*. Thus, *HMC03818* could cause a more severe fertility phenotype than *HMC04063* because its shRNA targets both genes. The absence of mature oocytes when using *MVD1* suggests that *CG18787* is required in early germline development. Both *HMC03818* did not cause lethality when crossed to *Tub:GAL4*, indicating *CG18787* may not be required for mitosis.

Mature oocytes expressing *HMC03818* had severe karyosome separation when compared to wildtype oocytes (54%, n=24, two examples in Figure 4E, 4F). This severe phenotype is consistent with a defect in sister chromatid cohesion during meiosis. The separation of sister kinetochores is a phenotype commonly associated with a loss of centromere cohesion [49]. Therefore, we determined the frequency of sister kinetochore separation in *HMC03818/ matα* oocytes. A normal meiosis is expected to have 8 foci, one for each pair of sister centromeres, although usually less is observed due to clustering of the centromeres. (avg = 6.9, Figure 4G). In contrast, *HMC03818* RNAi oocytes had a significant elevated frequency of sister kinetochore separation (avg = 8.8, p = 0.0185, Figure 4G). These results are consistent with the hypothesis that *G18787* is required for sister chromatid cohesion.

The defects associated with the loss of *CG18787* and *CG18789* may only appear in Metaphase I oocytes. Early meiotic prophase oocytes from *HMC03818 / MVD1* females were dissected and stained for SC component C(3)G and DSB marker ᵧH2AV (Figure 5D), but no cytological defects were observed.

*CG18787* contains a nucleoporin domain and limited homology to human *NUP42*. This is significant because recent studies have found that upon nuclear envelope breakdown leading up to cellular division, many nucleoporins localize to kinetochores and serve alternative functions to aid in cellular division. An example of this is the nucleoporin *Elys*, which was found to function as a Protein Phosphatase 1 scaffold during M-phase exit and thus aids in the disassembly of kinetochores [50]. Based on these results, we believe that *CG18787* has a meiotic function involved in the regulation of sister chromatid cohesion.

## Conclusion

Sexual reproduction depends on two integrated processes, the faithful transmission of chromosomes during meiosis to yield viable gametes, and the development of a gamete capable of fertilization and supporting embryonic development. Our unbiased screen in *Drosophila* has likely identified genes required for either or both processes.

In this study, we leveraged gene expression profiles and functional annotation to identify novel meiotic genes followed by phenotyping validation in *Drosophila* RNAi lines. We identified 65 genes that displayed elevated levels of nondisjunction or loss of fertility when knocked down, showing that these genes are required in multiple phases of gametogenesis and meiosis. We screened 253 RNAi lines using two *GAL4* promoters, which resulted in a selection for genes required early in gametogenesis or late in oogenesis. None of our 65 meiosis candidate genes were found in the manually curated meiosis database, MeiosisOnline [30]. These results supported our approach for identifying novel meiosis and fertility genes.

At least two previous studies used RNAi to screen for genes related to the loss of fertility. First was a screen using a *GAL4* promoter with similar expression characteristics to *MVD1* and *matα* for maternal proteins that are phospho-regulated [51]. Among the 132 genes whose knockdown affected oogenesis (class 2 to class 6), three genes overlap our 1,119 ovary up-regulated genes, including one of our positive genes *CG11188* and two genes that we tested but showed normal phenotype (*CG18259 /// CG6961, CG4968*). Second was a screen using a *GAL4* similar to *MVD1* for genes required for germline stem cell maintenance [52]. Among the 366 positive genes, 3 overlaps with our positive results (*CG13096*, *CG8435*, and *CG30020*) and 2 overlaps with our negative results (*CG42307*, and *CG11398*). The lack of overlap of these previous studies with our positive genes substantiates the validity of our approach of identifying novel genes. In the future, we could continue novel gene discovery by integrating ovary up-regulated genes in FlyAtlas 1 and FlyAtlas 2 [25]. While there are differences in experimental design and data generation methods between the two databases, the majority of the FlyAtlas1 ovary up-regulated genes (72.7%) are also up-regulated in FlyAtlas2, including 80% of the positive genes.

The genes we discovered fall into a variety of functional classes. The use of the two *GAL4* lines to induce expression of the shRNA also provides useful temporal information. Some genes, such as *CG7357, CG15863, CG5877, CG30020, CG42232,* and *CG7357*, were sterile with *MVD1* but not *matα*, indicating they had a function only in early germline development. Conversely, genes such *CG17361, CG6951, CG6540*, and *CG4554* were sterile with *matα* but not *MVD1*, suggesting they only function late in oocyte development. Knockdown of some genes failed to make mature oocytes. This phenotype with *matα*, such as *CG3430, CG14174, CG16838, CG33217,* and *CG6540*, suggests a role in growth of the oocyte during the meiotic prophase arrest. We also identified several genes with a reduced fertility phenotype. This could indicate a partial mRNA knockdown or a function important but not essential for fertility.

Several genes, such as *CG6937, CG3407, CG3430, CG12259, CG10635, CG10344, CG6843,* and *CG34261* had significant nondisjunction phenotypes and thus appear to be required for meiosis. Although several genes involved in homolog pairing and recombination within *Drosophila* show little apparent sequence homology, there is strong genetic and structural conservation of meiosis across eukaryotes [53–56]. The functional similarity between human and fruit fly establishes the foundation that understanding of the *Drosophila* genome could provide a valuable source to yield insights into human gene functions that are not easily obtainable in mammals like humans or mice. For example, the positive gene *CG8915* is the homolog of a potential human meiotic gene *YTHDC2*, which plays a role in regulating meiosis [57–59].

An important outcome of this screen is the identification of genes with no known meiotic function that have homologs in higher eukaryotic systems. For example, 57 of the 65 genes have mouse homologs, and one-third of these genes have no known function in meiosis or reproduction. The high percentage of human/mouse homologs in our positive genes demonstrate this approach enriches for genes that are crucial for oocyte and embryo development and can uncover novel mechanisms for female infertility. This screen also uncovered nearly 20 genes that either function or have predicted function in RNA biology, a function intimately linked to high egg quality. Therefore, this study has opened new and critical areas related to meiosis and egg quality that should be explored.

## Acknowledgement

This work is supported by NIH grants R01HD091331 and R01GM101955.

## Data Availability Statement

The data is available in the manuscript and the associated Supplemental Tables.

## Supplemental Figures

**Figure S 1:**
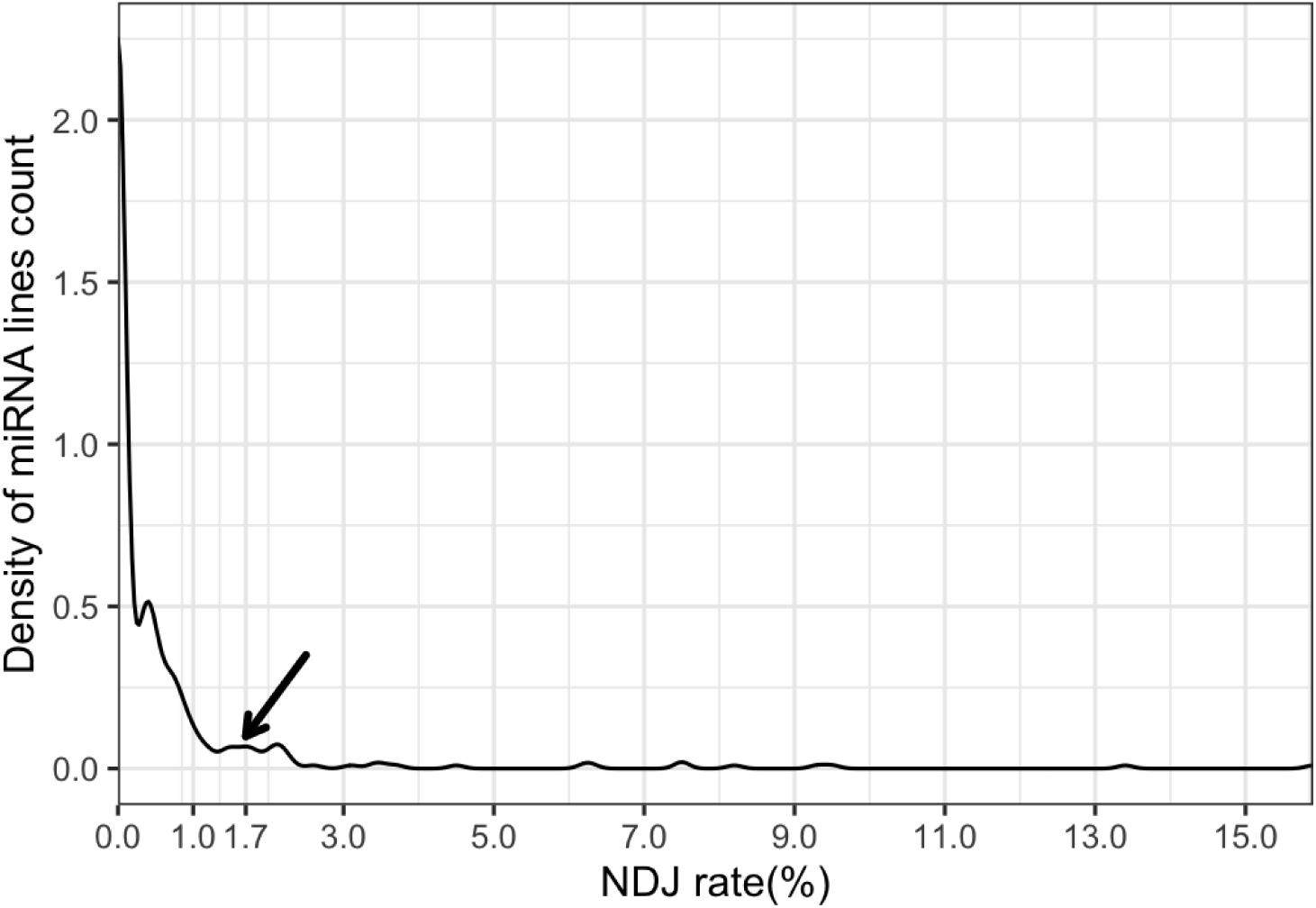
Distribution of RNAi knockdown non-disjunction results. All observed rates of nondisjunction were plotted to determine the frequency of occurrence. Elevated rates of nondisjunction were determined to be about 1.68% based on the distribution of frequencies.

## Supplemental Tables

**Table S1: Details of ovary up-regulated genes from FlyAtlas 1**

**Table S2: RNAi knockdown phenotypes**

**Table S3: Gene Ontology enrichment analysis for mouse and human orthologs of the positive genes**

**Table S4: Pathway enrichment analysis for mouse and human orthologs of the positive genes**

**Table S5: PPI network of positive genes**

